# H_2_S Prodrug, SG-1002, Protects Against Myocardial Oxidative Damage and Hypertrophy via Induction of Cystathionine β-Synthase and Antioxidant Proteins

**DOI:** 10.1101/2021.03.28.437435

**Authors:** Rahib K. Islam, Erinn Donnelly, Fokhrul Hossain, Jason D. Gardner, Kazi N. Islam

**Author notes:** Equal contribution (Rahib K. Islam, Erinn Donnelly). To whom correspondence should be addressed: Agricultural Research Development Program, College of Engineering, Science, Technology and Agriculture, Central State University, 1400 Brush Row Road, Wilberforce, OH.

## Abstract

Endogenously produced hydrogen sulfide (H_2_S) is critical for cardiovascular homeostasis. Therapeutic strategies aimed at increasing H_2_S levels have proven cardioprotective in models of acute myocardial infarction (MI) and heart failure (HF). The present study was undertaken to investigate the effects of a novel H_2_S prodrug, SG-1002, on stress induced hypertrophic signaling in murine HL-1 cardiac muscle cells. Treatment of HL-1 cells with SG-1002 under serum starvation without or with H_2_O_2_ increased the levels of H_2_S, H_2_S producing enzyme, cystathionine β-synthase (CBS) as well as antioxidant protein levels, such as super oxide dismutase1 (SOD1) and catalase and decreased oxidative stress. SG-1002 also decreased the expression of hypertrophic/HF protein markers such as atrial natriuretic peptide (ANP) and brain natriuretic peptide (BNP) in stressed HL-1 cells. Treatment with SG-1002 caused a significant induction of cell viability and a marked reduction of cellular cytotoxicity in HL-1 cells under serum starvation incubated or with H_2_O_2_. Experimental results of this study suggest that SG-1002 attenuates myocardial cellular oxidative damage and/or hypertrophic signaling via increasing H_2_S levels or H_2_S producing enzyme, CBS and antioxidant proteins.

## Introduction

The HL-1 cell line is derived from an AT-1 mouse atrial cardiomyocyte tumor and has several key advantages over other types of cardiac cells. It can be recovered from frozen and passaged indefinitely while still maintaining its differentiated biochemical and morphological features, as well as the ability to contract. Because of these properties, HL-1 cardiomyocytes can be used to model the effects of HF, hypertrophy and oxidative stress in vitro (1). The present study was undertaken to determine the effects of prodrug, SG-1002, a hydrogen sulfide (H_2_S) donor, on oxidative stress induced hypertrophic signaling in HL-1 cells. Sulfur has a critical role in protein structure/function and redox status/signaling in all living organisms. Although a sulfur containing molecule, H_2_S, is now recognized as a central player in physiology and pathobiology, the full scope and depth of sulfur metabolome’s impact on human health and longevity has been vastly underestimated and is only starting to be grasped. Since many pathological conditions have been related to abnormally low levels of H_2_S in blood and/or tissues, and are amenable to treatment by H_2_S supplementation, development of safe and efficacious H_2_S donors deserve to be undertaken with a sense of urgency; these prodrugs also hold the promise of becoming widely used for disease prevention and as antiaging agents. One such prodrug is an SG-1002, a precursor to a natural-occurring molecule, H_2_S, for which deficits have been shown to exist in a number of serious diseases including cardiovascular disease (CVD), type II diabetes, cancer, hypertension, etc.

Once thought of only as a toxic gas, H_2_S now belongs to a class of compounds known as gasotransmitters. H_2_S, along with carbon monoxide (CO) and nitric oxide (NO) are endogenously produced gases that are required for cardiovascular homeostasis (2). H_2_S is of particular interest as a possible agent to combat HF because of its ability to promote vasodilation and its anti-inflammatory (3), antioxidant (4) and anti-apoptotic (5) properties. Previous studies have shown that genetic overexpression of H_2_S producing enzymes protects against HF while deficiency exacerbates it (6). H_2_S is a potent antioxidant that can eliminate free radicals and prevent new reactive oxygen species (ROS) from forming, which are particularly detrimental in myocardial infarction/reperfusion (MI/R) injuries (4). Additionally, H_2_S has recently been implicated as a potential treatment against hypertrophic signaling (7), which is responsible for the pathological remodeling associated with HF.

H_2_S is made endogenously by three enzymes: cystathionine β-synthase (CBS), cystathionine γ-lyase (CSE), and 3-mercaptopyruvate sulfur transferase (3-MST). In mammals, CSE is predominately responsible for the manufacture of H_2_S in the cardiovascular system, while CBS is present in greater quantities in the central nervous system (8). 3-MST is responsible for manufacturing 90% of the H_2_S in the brain (8). A study by Islam et al. has shown that nitric oxide therapy protects cardiac function in MI murine model via induction of H_2_S producing enzymes (CSE and CBS)/H_2_S and antioxidants levels (9). In this study, we seek for a mechanism of the reduction in stress induced cardiomyocyte hypertrophic signaling through the use of an exogenous H_2_S donor, SG-1002. We also aimed to determine whether this prodrug could induce expression of CBS, CSE and 3-MST, which would allow cells to produce more endogenous H_2_S.

## Materials and Methods

### Cell Culture

Murine HL-1 cardiac muscle cells were maintained in serum containing (10%) media followed by serum starvation (1%) media for 24 hours. Serum starved/stressed cells were treated for 1 hour with either SG-1002 (Sulfagenix, Cleveland, OH), or H_2_O_2_ 500 (μM), or hypertrophic agonist such as endothelin-1 (ET-1) or in combination. Treated cells were analyzed for protein levels (by immunoblot), mRNA levels (by qRT-PCR), H_2_S levels, oxidative stress, and related assays.

### Real Time Quantitative Reverse Transcription Polymerase Chain Reaction (RTqRT-PCR)

RNA was isolated from the stressed HL-1 cells treated with or without one or two specific reagents for 1 hour in 1% serum containing media. 1 μg of RNA was used for the synthesis of cDNA. TaqMan primers for ANP, BNP, SOD-1, catalase, CSE, CBS, and 3-MST, were used to amplify q-PCR. 18s was used as a house keeping gene.

#### Measurement of H_2_S

The levels of H_2_S were determined in media samples by using gas chromatography chemiluminescence method as described previously (9).

#### Measurement of Advanced Oxidative Protein Products (AOPP)

All reagents or chemicals used in our experiments were purchased from Sigma-Aldrich. Advanced oxidative protein products AOPP Assay Kit was obtained from Abcam (ab242295). AOPP is a simple, reproducible, and consistent system for the detection of advanced oxidation protein products in plasma, lysates, and tissue homogenates. This kit includes a Chloramine standard and an AOPP Human Serum Albumin conjugate for use as a positive control. AOPP was determined using a spectrophotometric method. Samples were incubated with glacial acetic acid and the absorbance was read at 340 nm. Chloramine T with potassium iodide was used as calibrator.

#### Cell Proliferation assay

Cell viability was determined utilizing a Vybrant® MTT cell proliferation assay kit (Thermo Fisher Scientific). Cells were maintained on a 96-well tissue culture plate (Falcon, Corning, NY) in 10% serum and cultured for 24 hours in serum starvation (1%) prior to treatment. Cells were then pretreated with DMSO or SG-1002 for 30 minutes, followed by treatment with H_2_O_2_, ET-1 and/or SG-1002 for one hour. 10 μL of the 12 mM MTT stock solution was loaded into each well and the plate was incubated in the dark at 37°C for 2 hours. Following the incubation period, 175 μL of the media was removed, and 50 μL of DMSO was added to the remaining 25 μL in each well. The plate was then incubated at 37°C for 10 minutes and the optical density was determined using a spectrophotometer at 540 nm.

### Lactate Dehydrogenase (LDH) Cytotoxicity Assay

Treated HL-1 cellular cytotoxicity was determined by using the assay kits obtained from Thermo Scientific.

#### Immunoblot Assay

The polyclonal antibody for GPx-1/2 (sc-30147) was purchased from Santa Cruz Biotechnology, Inc. (Santa Cruz, CA USA). Monoclonal antibody for GAPDH (sc-32233) was also purchased from Santa Cruz Biotechnology, Inc. The polyclonal antibody for Catalase (H000000847-D01) was purchased from Abnova Corporation (Taipei City, Taiwan). Total protein samples were prepared from treated HL-1 cardiomyocytes using RIPA lysis buffer and quantified using BCA protein assay kits from Pierce, Inc. Samples were then separated via electrophoresis on a 4-20% Mini-PROTEAN TGX Precast Gel (Bio-Rad Laboratories, Inc.) and transferred to a 0.45 μM nitrocellulose membrane (Bio-Rad Laboratories, Inc.). Membranes were blocked for at least two hours with Odyssey Blocking Buffer (Li-Cor, Lincoln, NE), diluted 1:1 with 1x PBS followed by probing with primary and secondary antibodies.

#### Measurement of ROS

Cellular ROS were measured using a cellular ROS/superoxide detection assay kit (Abcam, Cambridge, UK). Cells were maintained on a 96-well tissue culture plate (Falcon, Corning, NY) in 10% serum and cultured for 24 hours in serum starvation (1%) prior to treatment. Cells were then pretreated with DMSO or SG-1002 for 30 minutes, followed by treatment with H_2_O_2_, and/or SG-1002 for one hour. After treatment, cells were loaded with 100 μL/well of ROS superoxide detection solution and incubated for one hour in the dark at 37°C. The plate was then read from the bottom at excitation 485/20 and emission 528/20.

## Results

### Time course and dose responses of SG-1002 on H_2_S production

HL-1 cardiac muscle cells were incubated either with different concentrations of SG-1002 for 1 hour (Fig. 1A) or with 10 μM of SG-1002 for several time points (Fig. 1B) in 1% serum containing media followed by the measurement of H_2_S levels in culture media. As can be seen 10 and 20 μM SG-1002 was able to significantly increase H_2_S levels in HL-1 cells (Fig 1A). It was also observed that treatment with 10 μM of SG-1002 produced more H_2_S in 1 hour as compared with other time points. This result suggests that SG-1002 induces the production H_2_S.

**Figure 1.**
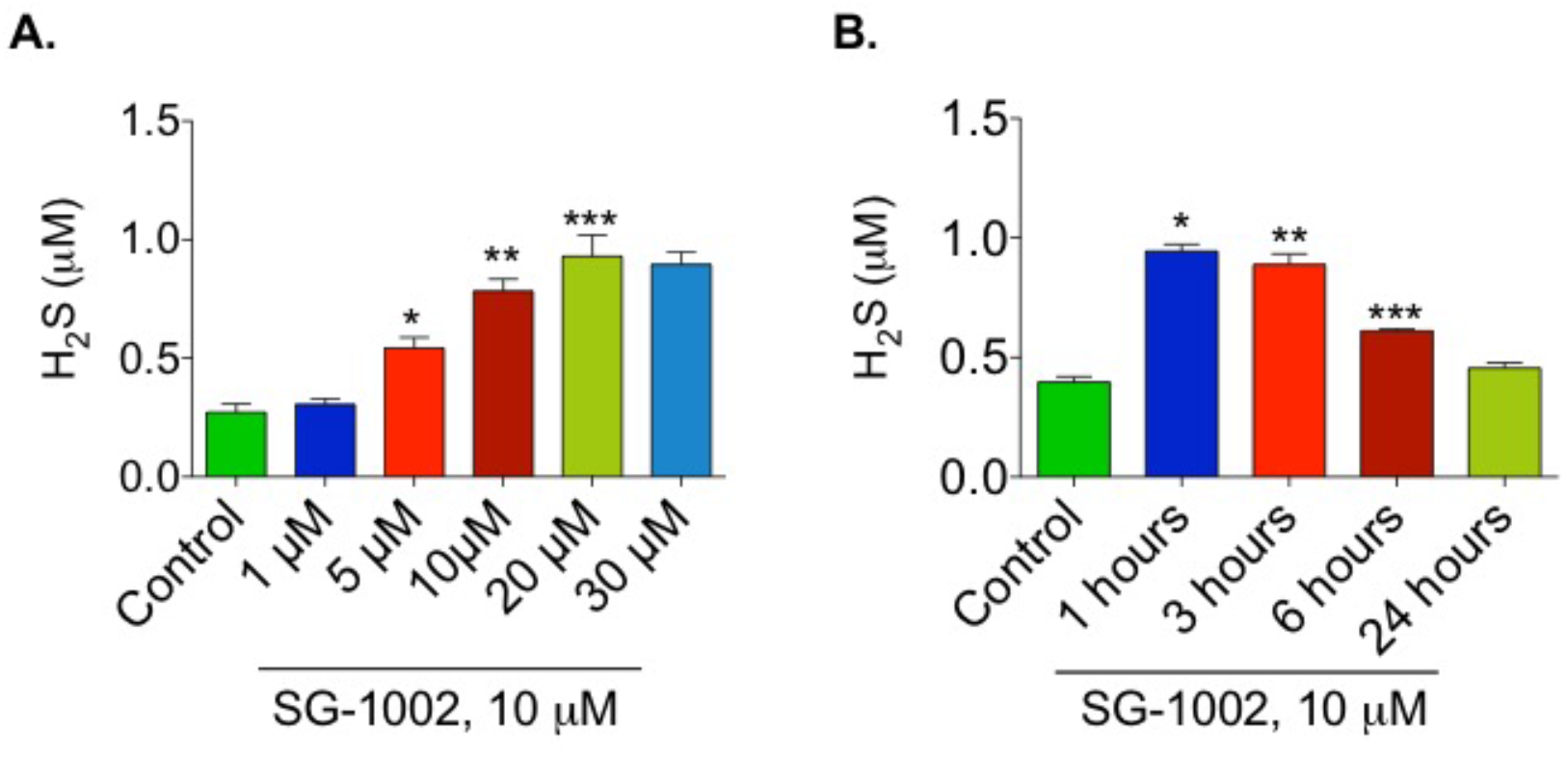
Time course and dose responses of SG-1002 on H_2_S production. HL-1 cardiac muscle cells were incubated either with different concentration of SG-1002 for 1 hour (A) or with 10 μM of SG-1002 for several time points (B) in 1% serum containing media followed by the measurement of H_2_S levels in culture media. *, **, and ***. P<0.05 versus (vs) control (n=4).

### Induction of CBS by SG-1002 in Cultured HL-1 Cardiac Muscle Cells

Treatment with SG-1002 significantly increased (p<0.05) the mRNA expression of CBS in HL-1 cardiac muscle cells cultured for 1 hour in 1% serum, as compared to the control. mRNA expression of CSE and 3-MST were also slightly up-regulated compared to the control, though not significantly (Fig. 2A, B, and C).

**Figure 2.**
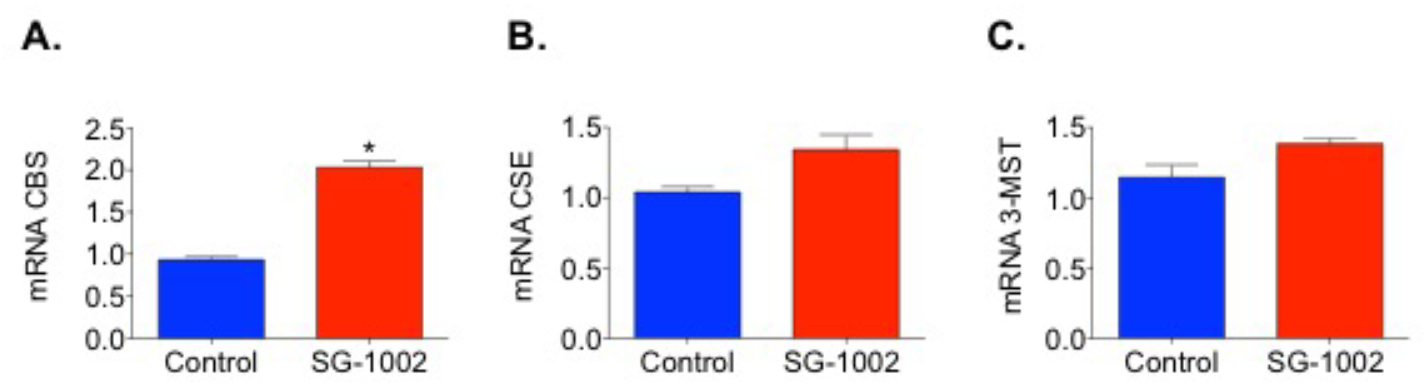
Effects of SG-1002 on mRNA Expression of CBS, CSE, and 3-MST in HL-1 cells. HL-1 cells were cultured with or without 10 μM of SG-1002 for 1 hour in 1% serum containing media. cDNA was prepared from RNA obtained from cultured HL-1 cells followed by analysis of mRNA of CBS (A), CSE (B), and 3-MST (C) using TaqMan PCR assay system. The experiment was repeated at least 3 times to verify the reproducibility. *, p<0.05 vs control, (A), (*t* test) (n=4).

### Effects of SG-1002 on the protein levels of CBS, CSE and 3-MST in HL-1 Cells

HL-1 cells were treated with SG-1002 (10 μM) or DMSO (control) for 1 hour following 24 hours of 1% serum starvation. Total protein was extracted from these cells and levels of CBS, CSE, and 3-MST were determined by immunoblot analysis. GAPDH was used as a loading control. CBS was significantly increased (p<0.05) in the SG-1002 treated cells as compared with control (Fig. 3A and B). CSE and 3-MST were also slightly increased in the SG-1002 treated cells, though not significantly (Fig. 3A, C, and D).

**Figure 3.**
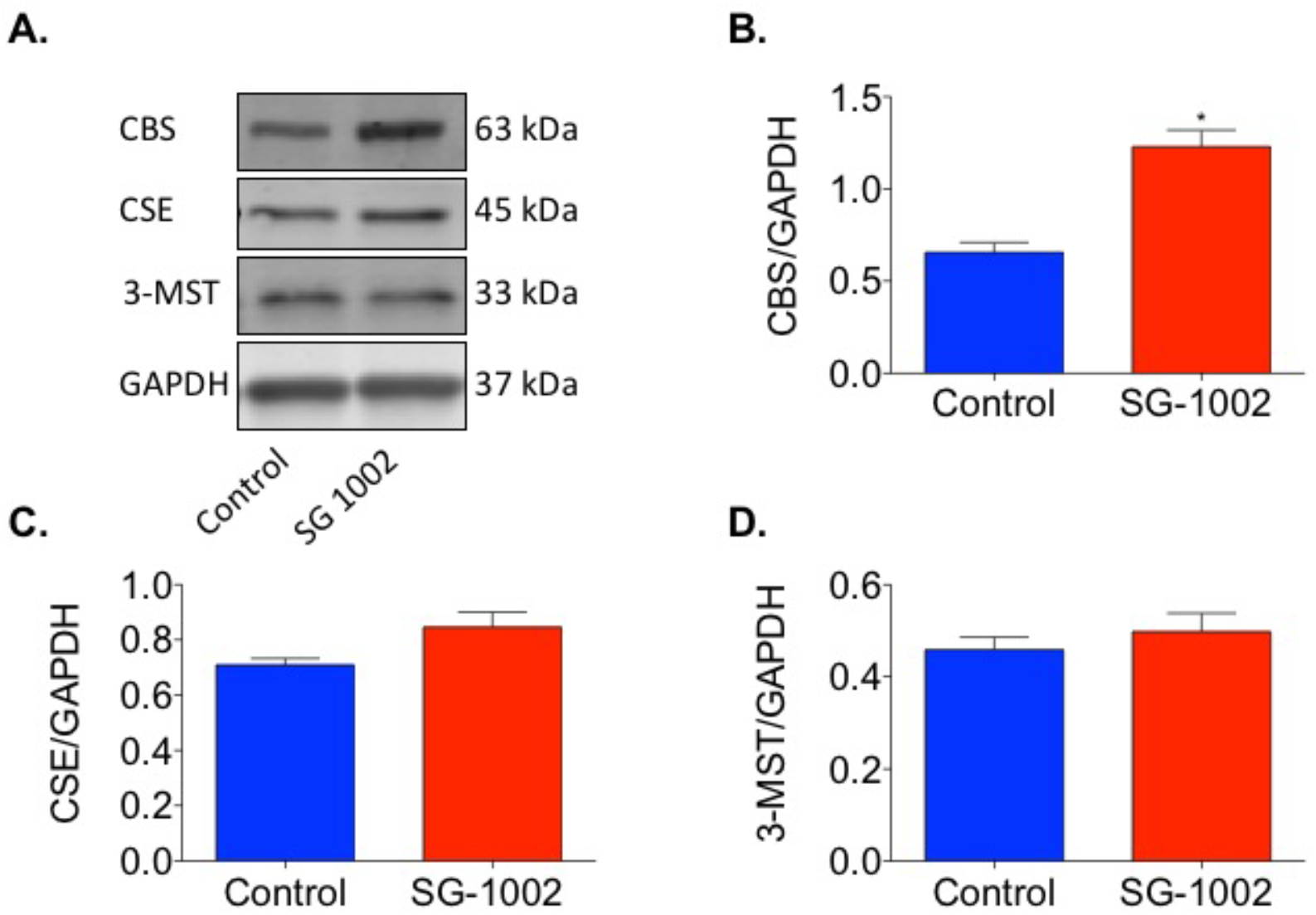
The effects of SG-1002 on the levels of CBS, CSE, and 3-MST in HL-1 cells. HL-1 cells were cultured with or without 10 μM of SG-1002 for 1 hour in 1% serum containing media followed protein extraction. CBS, CSE, and 3-MST levels were determined by using immunoblot analysis. (A) represents blots for CBS, CSE, and 3-MST and GAPDH. (B), (C), and (D) represent the quantitation of blots in (A). *, p<0.05 versus control, (B), (*t* test) (n=4).

### Inhibition of oxidative stress by SG-1002 in cultured HL-1 cardiac muscle cells

Hl-1 cells treated with SG-1002 had significantly lower levels of oxidative stress when compared to the controls. AOPP and ROS levels were measured in 4 treated groups of HL-1 cells following 24 hours of serum starvation: DMSO vehicle control, 10 μM SG-1002, DMSO and 500 μM H_2_O_2_ (to further stress the cells), or SG-1002 and 500 μM H_2_O_2_. The cells treated with SG-1002 alone had the lowest levels of oxidative stress (AOPP and ROS), followed by the DMSO-treated control group (Fig. 4A and B). The H_2_O_2_ group had the highest levels of oxidative stress and SG-1002 was able to antagonize the effects of H_2_O_2_ (Fig. 4A and B).

**Figure 4.**
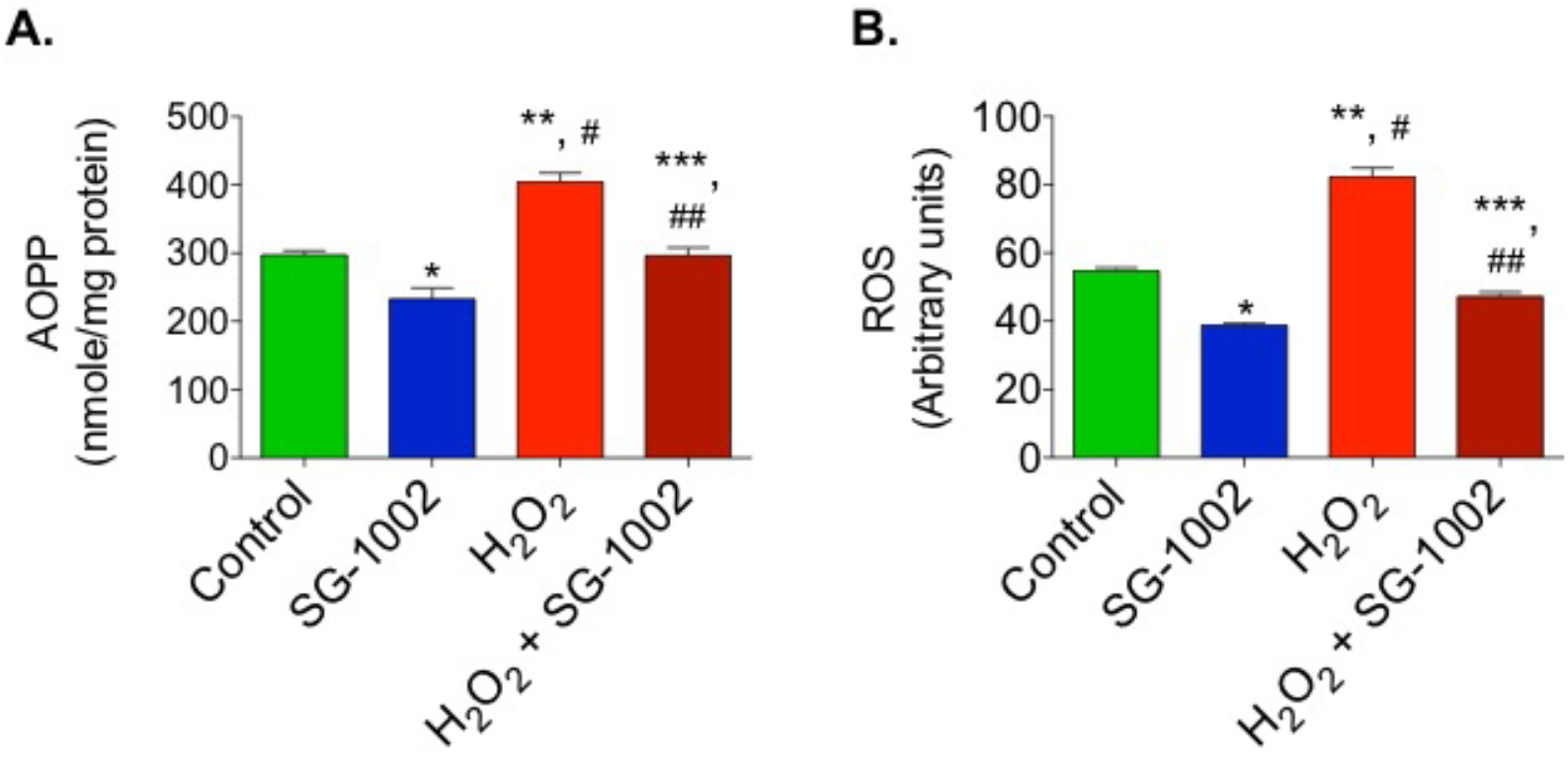
Reduction of oxidative stress in HL-1 cells by SG-1002. Oxidative stress was measured by determining the levels of AOPP (A) and ROS (B) in cultured HL-1 cells after incubation without or with either 10 μM of SG-1002 or 500 μM of H_2_O_2_ or in combination in 1% serum containing media. The experiment was repeated at least 3 times to verify the reproducibility. *, and **, p<0.05 vs control; ##, p<0.05 H_2_O_2_ VS SG-1002 + H_2_O_2_ (Anova).

### Effects of SG-1002 on mRNA expression of SOD1 and catalase in HL-1 cells

It was then of interest to determine the levels of antioxidative enzyme genes expression in HL-1 cells treated without or with either 10 μM of SG-1002, or H_2_O_2_ (500 μM) and or in combination for 1 hour in 1% serum containing media. cDNA was prepared from RNA obtained from cultured HL-1 cells followed by analysis of mRNA of SOD1 (Fig. 5A) and catalase (Fig. 5B) using TaqMan PCR assay system. Treatment of HL-1 cells with SG-1002 significantly increased mRNA expression of SOD1 and catalase. Furthermore, SG-1002 also increased the expression of both enzymes even under oxidative stress induced by H_2_O_2_ (Fig. 5A and B). This data suggests that SG-1002 protects HL-1 cells from oxidative damage via inducing antioxidant proteins.

**Figure 5.**
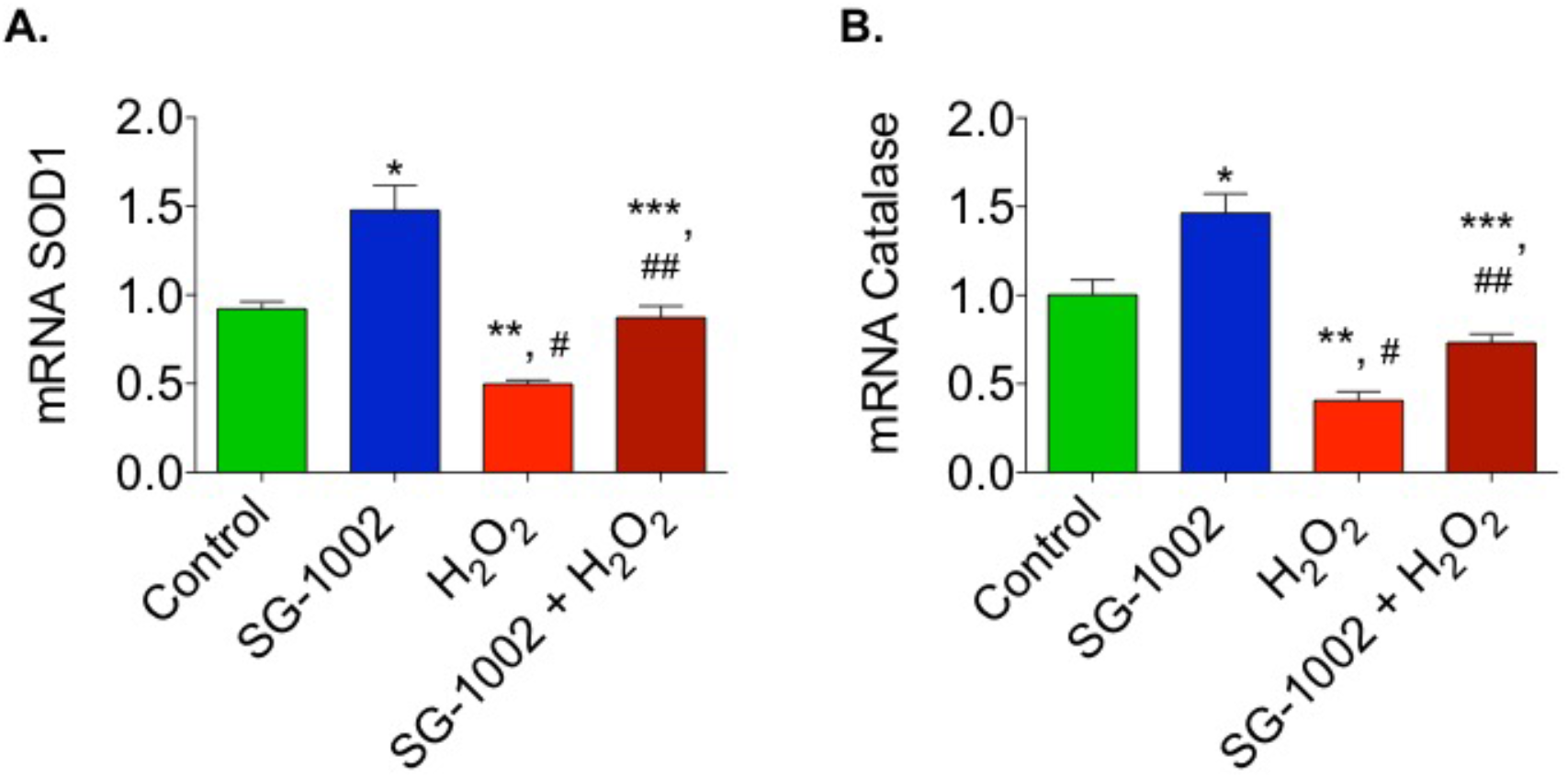
Effects of SG-1002 on mRNA expression of SOD1 and catalase in HL-1 cells. HL-1 cells were incubated either with 10 μM of SG-1002, or H_2_O_2_ (500 μM) and or in combination for 1 hour in 1% serum containing media. cDNA was prepared from RNA obtained from cultured HL-1 cells followed by analysis of mRNA of SOD1 (A) and catalase (B) using TaqMan PCR assay system. * and **, p<0.05 vs control; ##, p<0.05 H_2_O_2_ VS SG-1002 + H_2_O_2_; ***, p<0.05 SG 1002 vs SG 1002 + H_2_O_2_ (Anova), (n=4).

### Effects of SG-1002 on H_2_S production and CBS mRNA expression when cells were cultured under stress

It was also of interest to examine whether SG-1002 is able to induce production of H_2_S and H_2_S producing enzyme CBS, when cells were treated with H_2_O_2_. As can be seen here under oxidative stress SG-1002 also significantly induced the production of H_2_S as well as CBS mRNA in HL-1 cells treated with SG-1002 plus H_2_O_2_ as compared with H_2_O_2_ alone (Fig. 6A and B).

**Figure 6.**
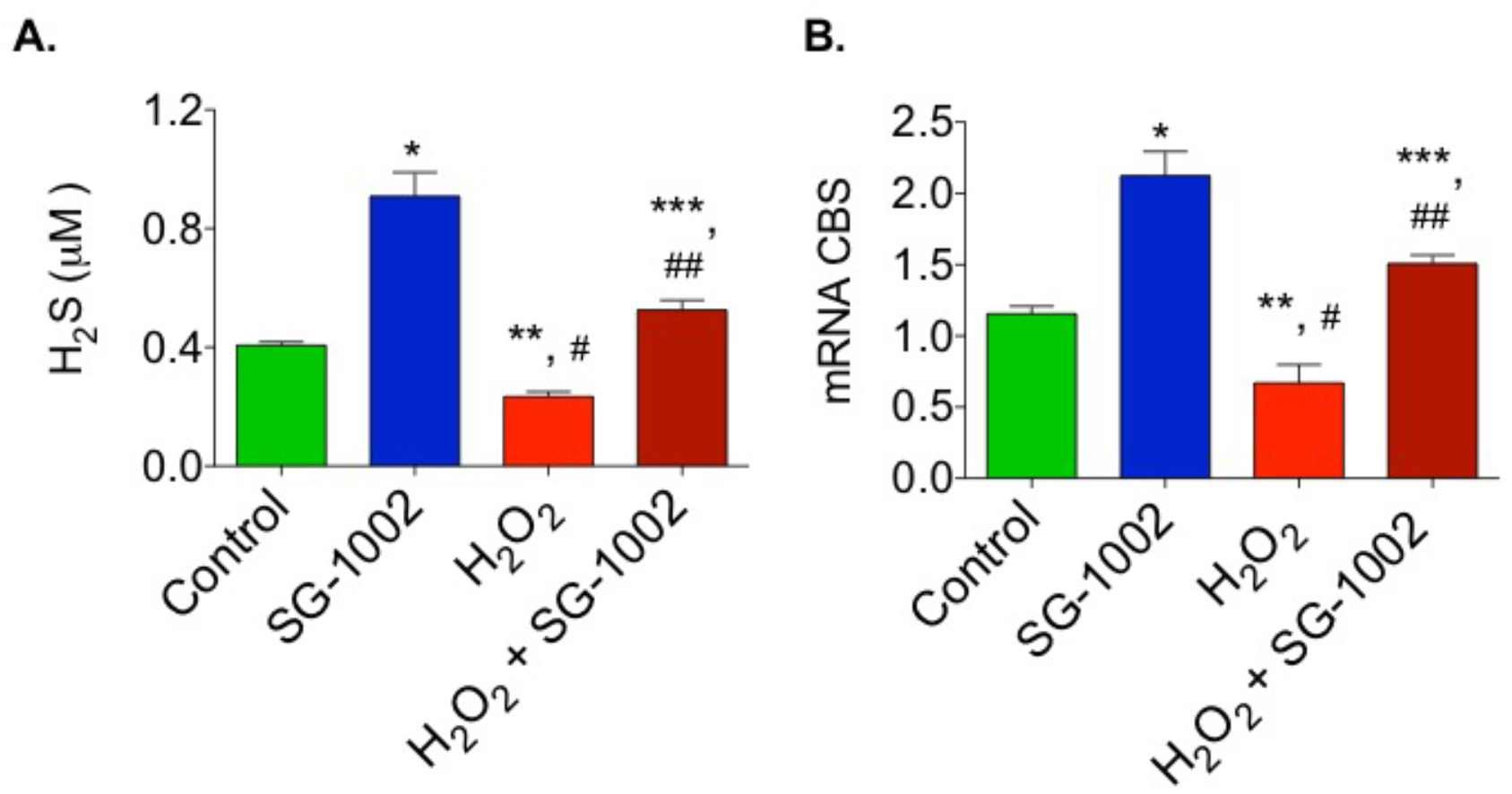
Effects of SG-1002 on H_2_S production and CBS mRNA expression when cells were cultured under stress. HL-1 cells were incubated either with 10 μM of SG-1002, or H_2_O_2_ (500 μM) and or in combination for 1 hour in 1% serum containing media followed by the measurement of H_2_S levels in culture media (A) and analysis of CBS mRNA (B) using TaqMan PCR assay system. *, and **, p<0.05 vs control; ##, p<0.05 H_2_O_2_ vs SG-1002 + H_2_O_2_; ***, p<0.05 SG-1002 vs SG 1002 + H_2_O_2_ (Anova) (n=3).

### Effects of SG-1002 on the expression of ANP and BNP in HL-1 cells

It is well known the expression of both ANP and BNP are elevated during hypertrophy or HF. Therefore, it was of interest to determine the levels of these gene expression in H_2_O_2_ and ET-1-treated HL1 cells. As can be seen in figure 7, SG-1002 significantly inhibited H_2_O_2_ or ET-1 induced ANP (Fig. 7A) and BNP (Fig. 7B) in HL-1 cells. SG-1002 inhibition of hypertrophic genes expression in stressed HL-1 cells suggests that SG-1002 plays important roles in the protection of HL-1 cells from oxidative damage.

**Figure 7.**
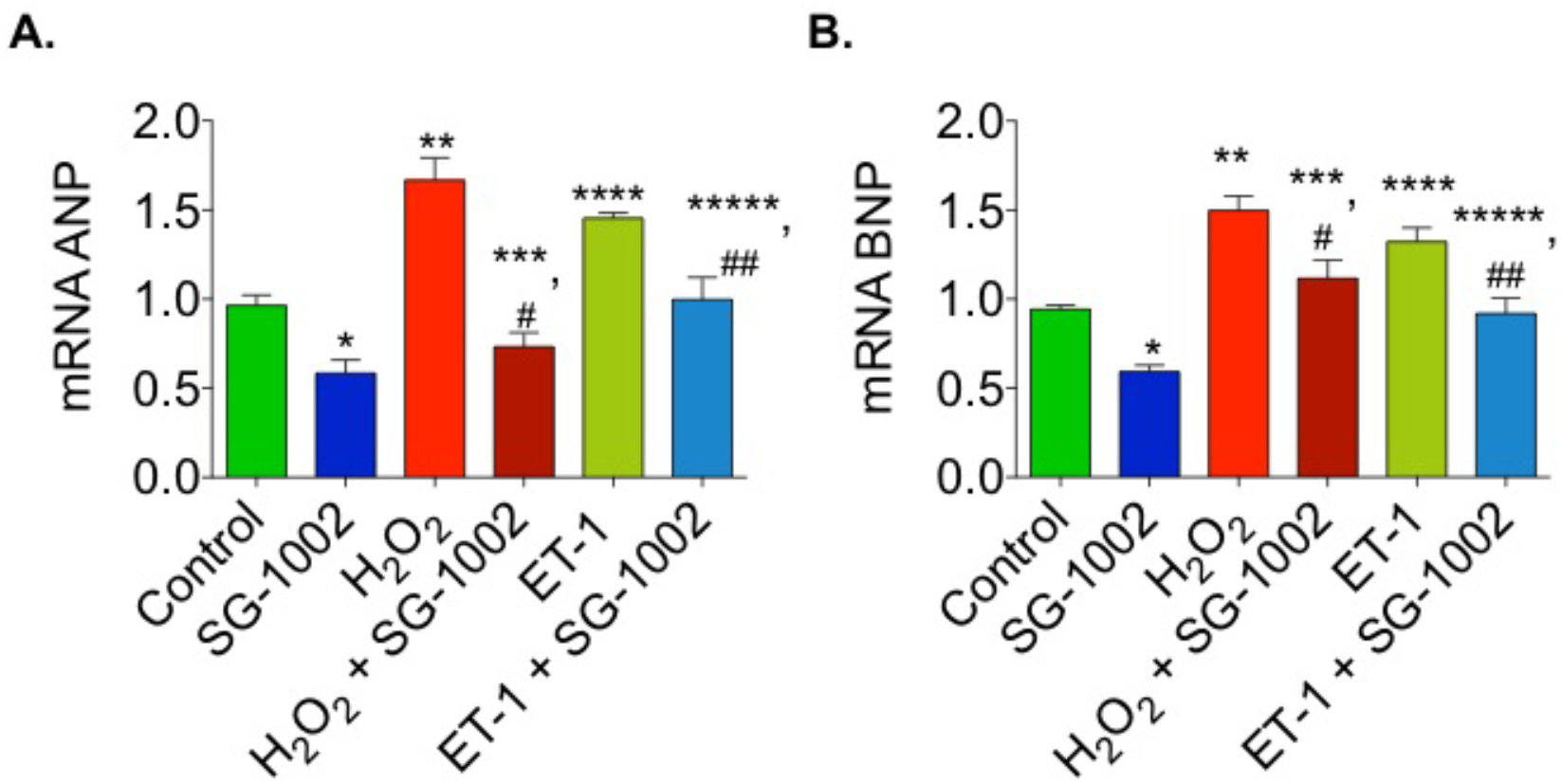
Effects of SG-1002 on the expression of ANP and BNP in HL-1 cells. Cells were cultured either with 10 μM of SG-1002 or H_2_O_2_ (500 μM) or ET-1 (10 nM) or in combination for 1 hour in 1% serum containing media. cDNA was prepared from RNA obtained from cultured HL-1 cells followed by analysis of mRNA of ANP (A) and BNP (B) using TaqMan PCR assay system. *, **, and ****, p<0.05 vs control; ^##^, p<0.05 ET-1 VS SG-1002 + ET1; ***, p<0.05 H_2_O_2_ VS SG-1002 + H_2_O_2_; (Anova) (n=4).

### SG-1002 decreases oxidative stress-induced cellular death and cytotoxicity in HL-1 cardiac muscle cells

Next, the effects of SG-1002 on cells viability and cytotoxicity were measured using HL1 cells treated without or with either 10 μM of SG-1002, or H_2_O_2_ (500 μM) and or in combination for 1 hour in 1% serum containing media. Cell viability (Fig. 8A) and cytotoxicity (Fig. 8B) were determined by using MTT and LDH cytotoxicity assays, respectively.

**Figure 8.**
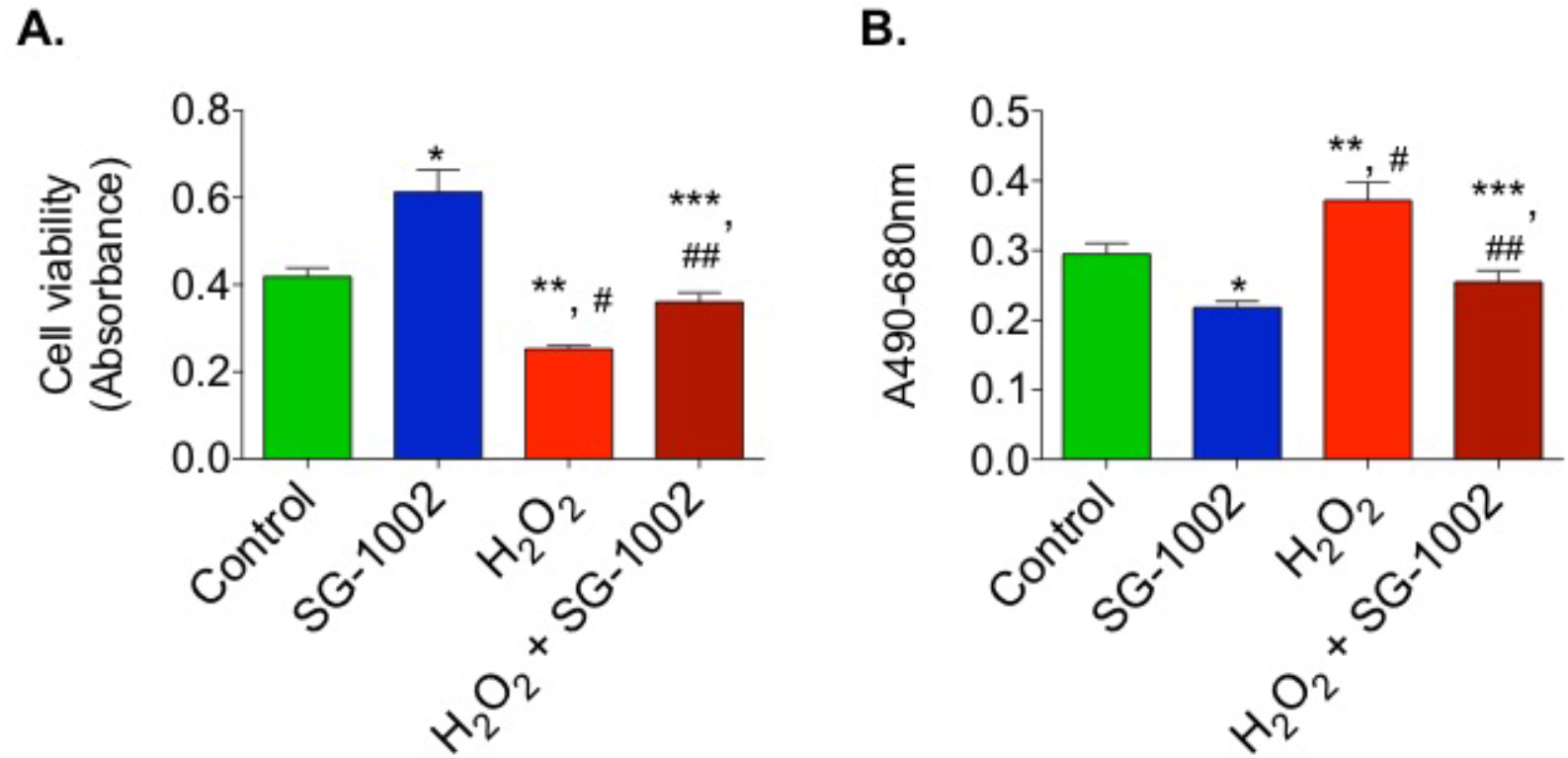
SG-1002 decreases oxidative stress-induced cellular death and cytotoxicity in HL-1 cardiac muscle cells. HL1 cells were incubated with either 10 μM of SG-1002, or H_2_O_2_ (500 μM) and or in combination for 1 hour in 1% serum containing media. Cell viability (A) and cytotoxicity (B) were determined by using MTT and LDH cytotoxicity assays, respectively. * and **, p<0.05 vs control; ##, p<0.05 H_2_O_2_ vs SG-1002 + H_2_O_2_; ***, p<0.05 SG-1002 vs SG 1002 + H_2_O_2_ (Anova) (n=3).

Treatment with SG-1002 alone markedly improves cell viability when compared with controls. Additionally, when the cells were further stressed with 500 μM H_2_O_2_, treatment with SG-1002 was able to significantly (p<0.05) improve cell viability as determined by MTT assay (Fig. 8A). Cytotoxicity was measured by using lactate dehydrogenase (LDH) cytotoxicity assay (Fig. 8B). Treatment with SG-1002 significantly lowered levels of LDH when compared to controls when HL-1 cells were under oxidative stress (Fig. 8B).

Data presented above indicate that treatment of HL-1 cells with SG-1002 induces H_2_S producing enzymes resulting in increased production of H_2_S. Increased levels of H_2_S activates nuclear factor erythroid 2-related factor 2 (Nrf2) which causes causing the elevation of antioxidant gene expression/antioxidants levels resulting in protection of cardiac cell from damage leading to the inhibition myocardial hypertrophy/HF (Fig. 9). The present data clearly demonstrate that SG-1002 attenuates myocardial cellular oxidative damage and/or hypertrophic signaling via increasing H_2_S levels/H_2_S producing enzyme CBS and upregulating antioxidant proteins.

**Figure 9.**
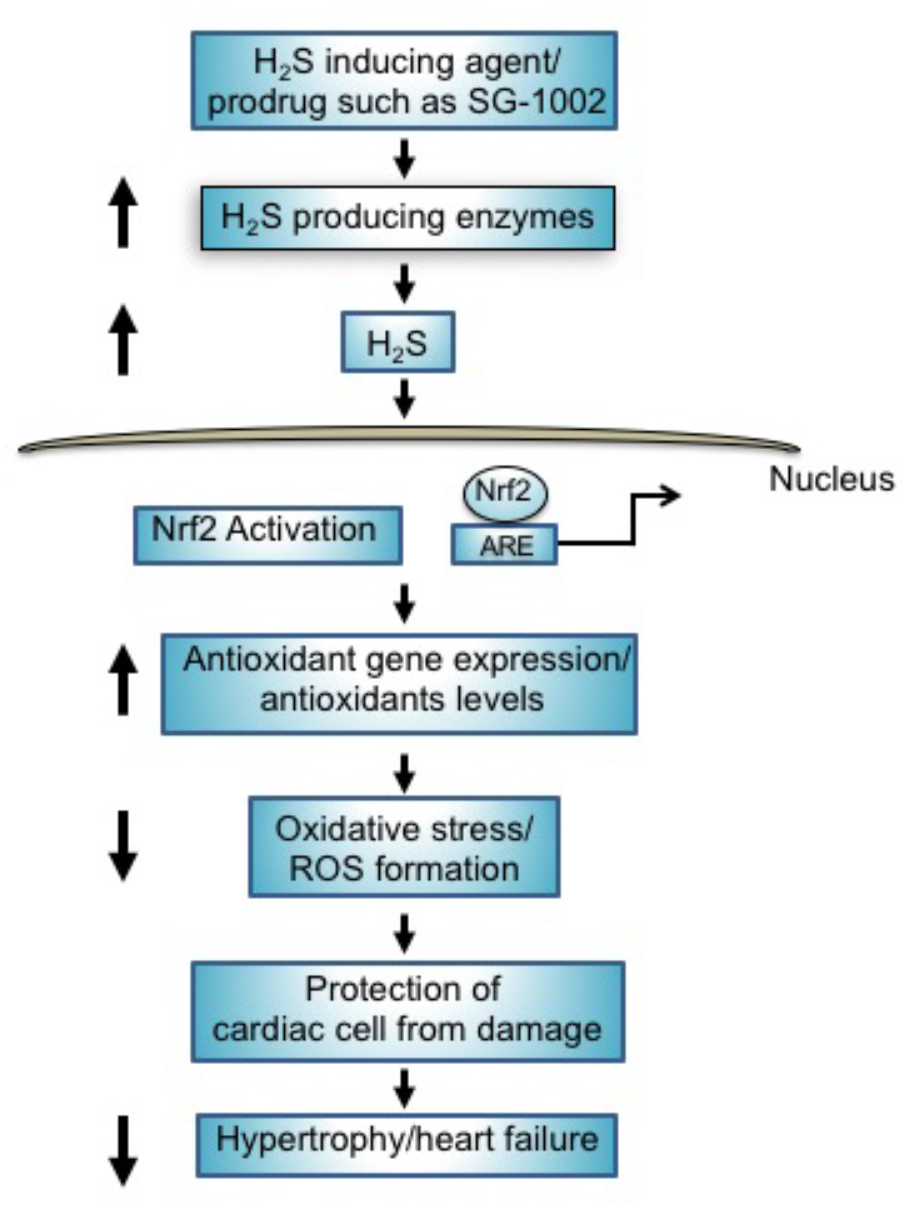
Mechanisms of protection of cardiac cells by H_2_S producing agents such as prodrug, SG-1002. Treatment of SG-1002 induces H_2_S producing enzymes resulting in increased production of H_2_S. Increased levels of H_2_S activates nuclear Nrf2 as well as causing elevation of antioxidant gene expression/antioxidants levels resulting in protection of cardiac cell from damage leading to the inhibition hypertrophy/HF.

## Discussion

Mammalian homeostasis is maintained through a plethora of signaling chemical species that regulate the function of cells, tissues, and organs; among them, only NO, CO, and H_2_S-the so called gasotransmitters-are endogenous diatomic or triatomic molecules that are capable of freely diffusing across cell membrane (10). These three small molecules play critical roles in both health and disease, as evidenced by the fact that most cells in our body are endowed with the enzymes required to produce them. These gasotransmitters usually operate in a concentrated and cooperative manner, and significant alterations in tissue concentration of any one of them either has detrimental physiological consequences or reflects a disease state. However, many data support the notion that limited bioavailability of H_2_S, NO (nitric oxide), or CO (carbon monoxide) may be counteracted by exogenously supplied H_2_S-partially through its action on endothelial NO synthase (11) and Nrf2 (12), respectively, but also by other means (12). H_2_S through Nrf2 (12), (13) is capable of transactivating more than 200 cytoprotective genes, thereby upregulating transcription of multiple antioxidant enzymes, phase II detoxifying enzymes, enzymes that catalyze the synthesis and regeneration of GSH, enzymes in charge of regulating NADPH regeneration, mitochondrial bioenergetics, and lipid metabolism. In our current study, we observed that SG-1002 increased levels of SOD and catalase in stressed HL1 cells via inducing H_2_S producing enzymes.

Many correlations have been established between low levels of H_2_S in blood or tissue and the onset of disease states related to oxidative cell damage, chronic inflammation, (14) immune dysfunction (14), (15), endoplasmic reticulum (ER) stress (16), (17) dysregulation of mitochondrial bioenergetics, (18) and hyperproliferation of cells or viruses (19); such correlations suggest the existence of causal links that are being vigorously scrutinized. Moreover, in some instances, an inverse relationship between disease progression and H_2_S level in blood and/or tissues has been established (20), (21), (22). Pathological conditions associated with so-called “H_2_S-poor” disease states are amenable to correction by H_2_S donors include (23), (24) aging, ischemia, cardiac hypertrophy, HF, liver disease (cirrhosis, steatosis), hypertension, atherosclerosis, endothelial dysfunction, diabetic complications, preeclampsia, Alzheimer’s disease (AD), and Huntington’s disease (HD).

DATS (diallyl trisulfide), DBTS (dibutenyl trisulfide), TC-2153 (benzopentathiepin 8-trifluoromethylv-1,2,3,4,5-benzopentathiepin-6-amine hydrochloride), and SG-1002 have the potential to become safe and effective pharmacological therapeutic agents that collectively will prove to be invaluable in humanity’s fight against the ravages of hundreds of disease conditions related to oxidative stress and cellular damage inflicted by ROS: These conditions include most aging related diseases. In the case of SG-1002, this potential beneficial effect is exclusively based on the fact that it is an H_2_S prodrug, whereas the other donors elicit pharmacologic effects that are only partially mediated by H_2_S. To realize the therapeutic potential of these four agents, it will be necessary to invest considerable resources to carry out the required clinical trials, since to the best of our knowledge only in one case (SG-1002) has safety been demonstrated in a formal Phase 1 clinical study.

Many previous studies have established that oxidative stress/ROS exacerbate HF while antioxidants protect against it (25). ROS is an important by-product of many cellular processes. Certain ROS, such as hydrogen peroxide (H_2_O_2_), act as signaling molecules (26) for the immune system and helps recruit white blood cells to initiate healing to damaged tissues (27). They are usually eliminated via interactions with superoxide dismutase, catalase, glutathione peroxidase, and peroxiredoxins (28). However, in many chronic conditions, more ROS are produced than can be eliminated, causing an imbalance between the antioxidant and oxidant systems. This imbalance is toxic and causes damage to many organelles including the mitochondria. In the setting of HF, this imbalance can cause changes in normal autophagy pathways (29), worsening arteriosclerosis (30), and persistent low levels of systemic inflammation (31).

In addition to the inflammation and oxidative stress, much of the pathology associated with heart disease comes from HF-induced hypertrophy and cardiac remodeling. Recent studies suggest that changes in the autophagy pathways may play a role in pathological remodeling (32). Reduced autophagy following an ischemia/reperfusion injury is associated with an increase in ROS and damage to the mitochondria, ultimately resulting in cell death (32).

Several mechanisms of action have been proposed during oxidative stress; superoxide molecules break down endothelium-derived NO, forming peroxynitrite, which, in addition to reducing the bioavailability of NO, causes DNA damage and oxidizes an eNOS cofactor (33). This process can be attenuated by the introduction of antioxidant compounds, such as sodium hydrosulfide (NaHS), which was found to inhibit the formation of NADPH oxidase derived superoxide (33). In a previous study it has been shown that CSE K/O mice had increased hypertension and decreased vasorelaxation compared to wild type animals (34). Another study found that mice that had undergone myocardial infarction/reperfusion (MI/R) had smaller infarct sizes and improved LV function when treated with a H_2_S donor (35). Our present study similarly found that treatment of stressed HL1 cells with H_2_S donor, SG-1002 increased levels of the H_2_S-producing enzyme CBS as well as levels of antioxidant proteins SOD1 and catalase leading to the protection of cells from damage. This suggests that SG-1002 works at least in part by increasing expression of antioxidant systems. Our previous study in HF murine model has shown that induction of H_2_S producing enzymes and or H_2_S causing the activation of antioxidant gene regulated transcription factor, Nrf2 resulting in increased antioxidant protection of cells from damage (9).

Reduction of oxidative stress results in inhibition of apoptosis or cell death. Other studies have shown that treatment with exogenous H_2_S in the form of garlic extracts reduced cell death, apoptosis and oxidative stress in cardiomyocytes (36). What’s more, they found that cells received no benefit from these compounds when they were treated with inhibitors of endogenously produced NO and H_2_S, NG-nitro-L-arginine methyl ester (LNAME) and propargylglycine (PAG), respectively(36), suggesting that sulfide donors can increase endogenous production of H_2_S by up-regulating the enzymes that produce it. Furthermore, a study found that hydrogen sulfide protects the heart by, among other things, increasing activation of the sirtuin-1 pathway (37). This pathway is thought to mediate the antioxidant/oxidant systems and promote the expression of antioxidant genes under oxidative stress (38). H_2_S is a powerful antioxidant compound that is capable of scavenging free radicals in vitro (39). Similarly, we observed in our study when stressed cardiomyocytes were treated with SG-1002 they had lower levels of oxidative stress because of the induction H_2_S producing enzyme such as CBS. We also noted that the cells that were further stressed with H_2_O_2_ and treated with SG-1002 had much lower levels of oxidative stress compared to the H_2_O_2_ alone group, but higher levels than both the SG-1002 alone and control group, suggesting that high levels of pre-existing oxidative stress impair its effect.

Changes in normal cell death and apoptosis are known to occur in HF (40). While some studies point to apoptosis after MI/R being cardioprotective, others point to it being a detrimental to LV function. We determined the effects of SG-1002 on cell death and cytotoxicity in stressed HL1 cells. The assays we used include MTT and LDH cytotoxicity, both of which provide information about the amount but not the pathways that cause cell death and cytotoxicity. In order to determine which pro-survival pathways are being affected by treatment with SG-1002, future studies may investigate increases in genes and proteins related to improved survival such as beclin-1, or decreases in pro-apoptotic markers, such as bax and caspase-3. Other studies have already implicated abnormalities in alternative splicing in these and other apoptotic genes in cardiovascular disease (CVD) (41).

Another way sulfide donors are said to have cardioprotective effects is by attenuating hypertrophy. A previous study found that mice treated with 20 mg/kg/day of SG-1002 had less cardiac enlargement, preserved LV function and less fibrosis after TAC when compared to mice that received the vehicle (6). As it has been reported previously that the elevated levels of hypertrophic genes, ANP and BNP in HF murine models were markedly decreased by NO mediated production of H_2_S (9). Similarly, in our present cellular study we found that SG-1002 treatment significantly reduced expression of ANP and BNP. Interestingly, an antioxidant property of SG-1002 was observed when oxidative stress induced by H_2_O_2_ in HL-1 was antagonized by SG-1002. A study found that exogenous H_2_S not only prevented hypertrophy in neonatal rat cardiac ventricular myocytes (NRCMs), but also improved the viability of the hypertrophic cells, possibly by altering glucose metabolism (42).

Our findings are consentient with the existing literature regarding SG-1002’s effect on hypertrophy, apoptosis and oxidative stress. In the future, it would be beneficial to look at how SG-1002 affects other clinical markers of HF, particularly pro-fibrotic markers. Other studies have already found H_2_S can reduce organ fibrosis, though the mechanism of how it produces this effect is unknown (43). In addition, other studies have found that another possible mechanism of H_2_S’s protective effect is through the mitochondria. H_2_S/CSE has been shown to decrease methylation of mitochondrial transcription factor A (TFAM) (44), which regulates mtDNA copy number and mitochondrial biogenesis (45). H_2_S can also improve the function of mitochondria under hypoxic conditions. Under normal conditions, H_2_S can reduce ATP production by inhibiting cytochrome c oxidase (46). However, under conditions where oxygen concentrations are low (such as in I/R injuries), CSE can be translocated to mitochondria, where it produces H_2_S, which helps to preserve the generation of ATP (47). Furthermore, Elrod et al. found that mice subjected to MI/R injury and treated with an H_2_S donor at the time of reperfusion had preserved mitochondrial function and membrane integrity, as well as smaller infarcts, less fractional shortening and preserved ejection fraction when compared to vehicle-treated mice (35). With the evidence of mitochondrial involvement growing, future studies may benefit from examining the effects of SG1002 on mitochondrial gene expression and function.

A small (n=18) phase 1 clinical trial found SG1002 to be a safe and well-tolerated drug in both healthy controls and HF patients (48). SG-1002 is likely to emerge as the H_2_S donor of choice on accounts of its unique ability to efficiently generate H_2_S without byproducts and in a slow and sustained mode. Larger clinical studies are being designed to further test the safety and efficacy of SG-1002 for the treatment of HF/hypertrophy and CVD. Our study and others have shown that H_2_S prodrugs like SG-1002 increase the levels of H_2_S producing enzymes and antioxidants while decreasing hypertrophic gene expression and oxidative stress, making it a promising novel therapeutic drug.

## Footnotes

CBS, cystathionine β-synthase; CSE, cystathionine g-lyase; 3-MST, 3-mercaptopyruvate sulfur transferase; SOD 1, superoxide dismutase 1; HF, heart failure; AOPP, advanced oxidative protein products; ANP, atrial natriuretic peptide; BNP, brain natriuretic peptide; H_2_S, hydrogen sulfide; NO, nitric oxide; ROS, reactive oxygen species; ET-1, endothelin-1; LDH, Lactate Dehydrogenase; Nrf2, nuclear factor erythroid 2-related factor 2.

## References

1. William C. Claycomb Nal, Jr., Beverly S. Stallworth, Daniel B. Egeland, Joseph B. Delcarpio Ab, and Nicholas J. Izzo, Jr. HL-1 cells: A cardiac muscle cell line that contracts and retains phenotypic characteristics of the adult cardiomyocyte. Proc Natl Acad Sci USA. 1998;95:2979–84.

2. Adrienne L. King DJP, Shashi Bhushan, Hiroyuki Otsuka, Kazuhisa Kondo, Chad K. Nicholson, Jessica M. Bradley, Kazi N. Islam, John W. Calvert, Ya-Xiong Tao, Tammy R. Dugas, Eric E. Kelley, John W. Elrod, Paul L. Huang, Rui Wang and David J. Lefer. Hydrogen sulfide cytoprotective signaling is endothelial nitric oxide synthase-nitric oxide dependent. Proceedings of the National Academy of Sciences. 2014;111(8):3182–7.

3. Li L, Rossoni G, Sparatore A, Lee LC, Del Soldato P, Moore PK. Anti-inflammatory and gastrointestinal effects of a novel diclofenac derivative. Free radical biology & medicine. 2007;42(5):706–19.

4. Eloise Streeter HHNaJLH. Hydrogen sulfide as a vasculoprotective factor. Medical Gas Research. 2013;3(9).

5. Kamat PK, Kalani A, Tyagi SC, Tyagi N. Hydrogen Sulfide Epigenetically Attenuates Homocysteine-Induced Mitochondrial Toxicity Mediated Through NMDA Receptor in Mouse Brain Endothelial (bEnd3) Cells. Journal of cellular physiology. 2015;230(2):378–94.

6. Kondo K, Bhushan S, King AL, Prabhu SD, Hamid T, Koenig S, et al. H(2)S protects against pressure overload-induced heart failure via upregulation of endothelial nitric oxide synthase. Circulation. 2013;127(10):1116–27.

7. Zhang L, Wang Y, Li Y, Li L, Xu S, Feng X, et al. Hydrogen Sulfide (H2S)-Releasing Compounds: Therapeutic Potential in Cardiovascular Diseases. Front Pharmacol. 2018;9:1066.

8. Ahmad A, Sattar MA, Rathore HA, Khan SA, Lazhari MI, Afzal S, et al. A critical review of pharmacological significance of Hydrogen Sulfide in hypertension. Indian journal of pharmacology. 2015;47(3):243–7.

9. Donnarumma E, Bhushan S, Bradley JM, Otsuka H, Donnelly EL, Lefer DJ, et al. Nitrite Therapy Ameliorates Myocardial Dysfunction via H2S and Nuclear Factor-Erythroid 2-Related Factor 2 (Nrf2)-Dependent Signaling in Chronic Heart Failure. J Am Heart Assoc. 2016;5(8).

10. Mathai JC, Missner A, Kugler P, Saparov SM, Zeidel ML, Lee JK, et al. No facilitator required for membrane transport of hydrogen sulfide. Proc Natl Acad Sci U S A. 2009;106(39):16633–8.

11. Szabo C. Gasotransmitters in cancer: from pathophysiology to experimental therapy. Nat Rev Drug Discov. 2016;15(3):185–203.

12. Hourihan JM, Kenna JG, Hayes JD. The gasotransmitter hydrogen sulfide induces nrf2-target genes by inactivating the keap1 ubiquitin ligase substrate adaptor through formation of a disulfide bond between cys-226 and cys-613. Antioxid Redox Signal. 2013;19(5):465–81.

13. Tonelli C, Chio IIC, Tuveson DA. Transcriptional Regulation by Nrf2. Antioxid Redox Signal. 2018;29(17):1727–45.

14. Li L, Moore PK. Could hydrogen sulfide be the next blockbuster treatment for inflammatory disease? Expert Rev Clin Pharmacol. 2013;6(6):593–5.

15. Miller TW, Wang EA, Gould S, Stein EV, Kaur S, Lim L, et al. Hydrogen sulfide is an endogenous potentiator of T cell activation. J Biol Chem. 2012;287(6):4211–21.

16. Kabil O, Vitvitsky V, Banerjee R. Sulfur as a signaling nutrient through hydrogen sulfide. Annu Rev Nutr. 2014;34:171–205.

17. Wu J, Pan W, Wang C, Dong H, Xing L, Hou J, et al. H2S attenuates endoplasmic reticulum stress in hypoxia-induced pulmonary artery hypertension. Biosci Rep. 2019;39(7).

18. Szczesny B, Modis K, Yanagi K, Coletta C, Le Trionnaire S, Perry A, et al. AP39, a novel mitochondria-targeted hydrogen sulfide donor, stimulates cellular bioenergetics, exerts cytoprotective effects and protects against the loss of mitochondrial DNA integrity in oxidatively stressed endothelial cells in vitro. Nitric Oxide. 2014;41:120–30.

19. Alshorafa AK, Guo Q, Zeng F, Chen M, Tan G, Tang Z, et al. Psoriasis is associated with low serum levels of hydrogen sulfide, a potential anti-inflammatory molecule. Tohoku J Exp Med. 2012;228(4):325–32.

20. Chen YH, Yao WZ, Geng B, Ding YL, Lu M, Zhao MW, et al. Endogenous hydrogen sulfide in patients with COPD. Chest. 2005;128(5):3205–11.

21. Jiang HL, Wu HC, Li ZL, Geng B, Tang CS. [Changes of the new gaseous transmitter H2S in patients with coronary heart disease]. Di Yi Jun Yi Da Xue Xue Bao. 2005;25(8):951–4.

22. Kovacic D, Glavnik N, Marinsek M, Zagozen P, Rovan K, Goslar T, et al. Total plasma sulfide in congestive heart failure. J Card Fail. 2012;18(7):541–8.

23. Szabo C, Papapetropoulos A. International Union of Basic and Clinical Pharmacology. CII: Pharmacological Modulation of H2S Levels: H2S Donors and H2S Biosynthesis Inhibitors. Pharmacol Rev. 2017;69(4):497–564.

24. Zivanovic J, Kouroussis E, Kohl JB, Adhikari B, Bursac B, Schott-Roux S, et al. Selective Persulfide Detection Reveals Evolutionarily Conserved Antiaging Effects of S-Sulfhydration. Cell Metab. 2019;30(6):1152–70 e13.

25. Rosenbaugh EG, Savalia KK, Manickam DS, Zimmerman MC. Antioxidant-based therapies for angiotensin II-associated cardiovascular diseases. American Journal of Physiology - Regulatory, Integrative and Comparative Physiology. 2013;304(11):R917–R28.

26. Veal EA, Day AM, Morgan BA. Hydrogen peroxide sensing and signaling. Molecular cell. 2007;26(1):1–14.

27. Niethammer P, Grabher C, Look AT, Mitchison TJ. A tissue-scale gradient of hydrogen peroxide mediates rapid wound detection in zebrafish. Nature. 2009;459(7249):996–9.

28. Islam KN, Takahashi M, Higashiyama S, Myint T, Uozumi N. Fragmentation of ceruloplasmin following non-enzymatic glycation reaction. J Biochem. 1995;118(5):1054–60.

29. Li B, Chi RF, Qin FZ, Guo XF. Distinct changes of myocyte autophagy during myocardial hypertrophy and heart failure: association with oxidative stress. Experimental physiology. 2016.

30. Libby P. Inflammation in Atherosclerosis. Nature. 2002;420:868–74.

31. Ledwaba L, Tavel JA, Khabo P, Maja P, Qin J, Sangweni P, et al. Pre-ART levels of inflammation and coagulation markers are strong predictors of death in a South African cohort with advanced HIV disease. PloS one. 2012;7(3):e24243.

32. Ma X, Liu H, Foyil SR, Godar RJ, Weinheimer CJ, Hill JA, et al. Impaired autophagosome clearance contributes to cardiomyocyte death in ischemia/reperfusion injury. Circulation. 2012;125(25):3170–81.

33. Al-Magableh MR, Kemp-Harper BK, Hart JL. Hydrogen sulfide treatment reduces blood pressure and oxidative stress in angiotensin II-induced hypertensive mice. Hypertension research: official journal of the Japanese Society of Hypertension. 2015;38(1):13–20.

34. Yang G, Wu L, Jiang B, Yang W, Qi J, Cao K, et al. H2S as a physiologic vasorelaxant: hypertension in mice with deletion of cystathionine gamma-lyase. Science. 2008;322(5901):587–90.

35. Elrod JW, Calvert JW, Morrison J, Doeller JE, Kraus DW, Tao L, et al. Hydrogen sulfide attenuates myocardial ischemia-reperfusion injury by preservation of mitochondrial function. Proc Natl Acad Sci U S A. 2007;104(39):15560–5.

36. Xavier Lieben Louis RM, Sijo Joseph Thandapilly, Liping Yu and Thomas Netticadan. Garlic extracts prevent oxidative stress, hypertrophy and apoptosis in cardiomyocytes: a role for nitric oxide and hydrogen sulfide. BMC Complementary & Alternative Medicine. 2012;12(140):10.

37. Wu D, Hu Q, Liu X, Pan L, Xiong Q, Zhu YZ. Hydrogen sulfide protects against apoptosis under oxidative stress through SIRT1 pathway in H9c2 cardiomyocytes. Nitric oxide: biology and chemistry / official journal of the Nitric Oxide Society. 2015;46:204–12.

38. Alcendor RR, Gao S, Zhai P, Zablocki D, Holle E, Yu X, et al. Sirt1 regulates aging and resistance to oxidative stress in the heart. Circulation research. 2007;100(10):1512–21.

39. Ashfaq Ahmad MZAS, Hassaan A. Rathore, Abdullah I. Hussain, Safia Akhtar Khan, Tabina Fatima, Sheryar Afzal, Nor A. Abdullah and Edward J. Johns. Antioxidant Activity and Free Radical Scavenging Capacity of L-Arginine and NaHS: A Comparative In Vitro Study. Acta Poloniae Pharmaceutica - Drug Research. 2015;72(2):245–52.

40. Roberta A. Gottlieb KOB, Robert A. Kioner, Bernard M Babior, and Robert L. Engler. Reperfusion Injury Induces Apoptosis in Rabbit Cardiomyoctes. The Journal of Clinical Investigation, Inc 1994;94:1621–8.

41. Dlamini Z, Tshidino SC, Hull R. Abnormalities in Alternative Splicing of Apoptotic Genes and Cardiovascular Diseases. International journal of molecular sciences. 2015;16(11):27171–90.

42. Liang M, Jin S, Wu DD, Wang MJ, Zhu YC. Hydrogen sulfide improves glucose metabolism and prevents hypertrophy in cardiomyocytes. Nitric oxide: biology and chemistry / official journal of the Nitric Oxide Society. 2015;46:114–22.

43. Zhang S, Pan C, Zhou F, Yuan Z, Wang H, Cui W, et al. Hydrogen Sulfide as a Potential Therapeutic Target in Fibrosis. Oxidative medicine and cellular longevity. 2015;2015:593407.

44. Li S, Yang G. Hydrogen Sulfide Maintains Mitochondrial DNA Replication via Demethylation of TFAM. Antioxid Redox Signal. 2015;23(7):630–42.

45. Clayton DA. Replication and Transcription of Vertebrate Mitochondrial DNA. Annu Rev Cell Biol. 1991;7:453–78.

46. Blackstone E, Morrison M, Roth MB. H2S induces a suspended animation-like state in mice. Science. 2005;308(5721):518.

47. Fu M, Zhang W, Wu L, Yang G, Li H, Wang R. Hydrogen sulfide (H2S) metabolism in mitochondria and its regulatory role in energy production. Proceedings of the National Academy of Sciences of the United States of America. 2012;109(8):2943–8.

48. Polhemus DJ, Li Z, Pattillo CB, Gojon G, Sr., Gojon G, Jr., Giordano T, et al. A novel hydrogen sulfide prodrug, SG1002, promotes hydrogen sulfide and nitric oxide bioavailability in heart failure patients. Cardiovascular therapeutics. 2015;33(4):216–26.

